# Characterization of the calmodulin-like protein family in *Chara braunii* and their conserved interaction with the calmodulin-binding transcription activator family

**DOI:** 10.1101/2024.06.06.597814

**Authors:** Kyle Symonds, Udo Wali, Liam Duff, Wayne A. Snedden

## Abstract

Calcium sensor proteins play important roles by detecting changes in intracellular calcium and relaying that information onto downstream targets through protein-protein interaction. Very little is known about calcium sensors from plant species that predate land colonization and the evolution of embryophytes. Here, we examined the genome of the multicellular algae, *Chara braunii*, for orthologs to the evolutionarily-conserved calcium sensor calmodulin (CaM), and for CaM-like proteins (CMLs). We identified one CaM and eight CML isoforms which rang in size from 16.4 to 21.3 kDa and are predicted to have between two to four calcium-binding (EF-hand) domains. Using recombinant protein, we tested whether CbCaM and CbCMLs1-7 possess biochemical properties of typical calcium sensors. CbCaM and the CbCMLs all displayed high-affinity calcium binding with estimated global *K*_D_ values in the physiological µM range. In response to calcium binding, CbCaM and the CbCMLs exhibited varying degrees of increase in exposed hydrophobicity, suggesting different calcium-induced conformational changes occur among isoforms. We found many examples of putative CaM targets encoded in the *C. braunii* genome and explored the ability of CbCaM and CbCMLs to interact *in planta* with a representative putative target, a *C. braunii* CaM-binding transcription factor (CbCAMTA1). CbCaM, CbCML2, and CbCML4 associated with the C-terminal region of CbCAMTA1. Collectively, our data support the hypothesis that complex calcium signaling and sensing networks involving CaM and CMLs evolved early in the green lineage. Similarly, it seems likely that calcium-mediated regulation of transcription occurs in *C. braunii* via CAMTAs and is an ancient trait predating embryophytic emergence.

**Highlights:** Although calmodulin (CaM) and calmodulin-like (CML) proteins are well studied in vascular plants, little is known about their orthologs in ancient lineages. We characterized CaM and CMLs from *Chara braunii*, and assessed their ability to bind a representative target protein, a calmodulin-binding transcription factor, CbCAMTA1.

## Introduction

The ability to detect and respond to environmental stimuli is a universal property among living organisms. Given their predominantly sessile nature, plants have evolved unique proteins and pathways to sense, interpret, and respond to environmental signals and information (Edel et al., 2017; Luan and Wang, 2021). A main player in plant cellular signaling is the secondary messenger, calcium. High concentrations of calcium in the cytoplasm are toxic, and thus calcium is actively transported out of the cytosol into the apoplast, vacuole, or the endoplasmic reticulum. Various biotic and abiotic stimuli trigger the opening of calcium channels where calcium moves rapidly down its electrochemical gradient into the cytosol and activates signal transduction pathways that help coordinate cellular responses. The spatial and temporal patterns of calcium influx are often complex, with repeating oscillations that have been proposed to encode information about the nature of the stimulus (Kudla et al., 2018; Tian et al., 2020). There are two main hypotheses on how fluxes of calcium might encode information; (i) the calcium signature hypothesis and (ii) the calcium threshold hypothesis. The former posits that specific calcium patterns encode information regarding the nature of the stress, similar to a Morse code, and therefore must be decoded and interpreted by cell components. The calcium threshold hypothesis proposes that concentration thresholds are the main factor affecting differential perception and response to calcium signals. There is supporting evidence for each of these current theories, and they are not mutually exclusive, nor do they preclude the importance and contribution of other second messengers (Behera et al., 2018).

In order to propagate information during signaling cascades, calcium signals must be perceived in the cytoplasm by calcium-sensor proteins. Plants possess three main families of calcium-sensors; Calcium-dependent Protein Kinases (CDPKs), Calcineurin-B-like (CBL) proteins and their associated CBL-interacting Protein Kinases (CIPKs), and the Calmodulin (CaM) and CaM-like (CML) protein families (DeFalco et al., 2009; Kudla et al., 2018). CaM is an essential calcium sensor protein and is ubiquitous across eukaryotic taxa as it is the fourth most evolutionarily conserved protein in Eukaryota (Copley et al., 1999). The model plant, Arabidopsis has seven *CaM* genes and an expanded family of 50 *CML* genes that are thought to have evolved from *CaM* (McCormack and Braam, 2003; Bender and Snedden 2013). Whereas CaMs are widespread, CMLs are unique to plants and some protists including brown and red algae (Bender and Snedden, 2013). Although the roles of most plant CMLs remain unclear, some have been demonstrated to function as calcium sensors during growth and development and in response to environmental stress (Li et al., 2023; Zeng et al., 2023). Like CaM, CML proteins are non-catalytic sensor-relay proteins, and their functions depend predominantly on the targets that they interact with. Whereas CaM has many well-studied targets, the list of known targets for Arabidopsis CMLs is much smaller but is expanding as research continues. For example, AtCML8 interacts with the receptor kinase, AtBRI1, and reduces AtBRI1 autophosphorylation activity in *E. coli* (Oh et al., 2012). Another example is the interaction of AtCML36 with the calcium-ATPase pump, ACA8, which increases its activity in yeast microsomes (Astegno et al., 2017). More recently, the paralogs AtCML13 and AtCML14 were found to interact with the unconventional CaM-binding IQ (Iso-Gln) domains in IQ67 Domain (IQD) proteins, CaM-binding transcription activators (CAMTAs), and myosin motor proteins (Teresinski et al., 2023). AtCML13 and AtCML14, along with CaM, function as myosin light chains and regulate myosin velocity *in vitro* (Symonds et al., 2024c). AtCML13 and AtCML14 are important in plant development (Symonds et al., 2024b) and also function in CAMTA-mediated transcriptional regulation under salinity stress (Hau et al., 2023). Additional examples of putative and demonstrated CML targets have been reported, and this list is expected to grow as more tools become available for high-throughput protein-protein interaction analyses. In general, it is clear that CMLs, alongside CDPKs, and CBLs/CIPKs function as important calcium sensors during information processing in plant cells.

Although a number of studies have explored the diversity, properties, and functions of CMLs in various plant species (La Verde et al., 2018; Wang et al., 2023), there are very few reports describing CMLs from taxa that predate land colonization. The recent completion of the *Chara braunii* genome provides the opportunity to explore the properties of the CaM/CML protein family from a multicellular algae (Nishiyama et al., 2018). *C. braunii* is thought to have a last common ancestor with higher plant species around 450 million years ago (Zhu et al., 2015). Moreover, there is a rich history of using *C. braunii* and *C. corallina* as model organisms in cell biology, dating back to the early development of the compound microscope by Robert Hooke in 1665 (Hooke, 1665; as cited in Nebenführ and Dixit, 2018). The large cell size (>5 cm in length; Nishiyama et al., 2018) of *C. braunii* facilitated foundational studies of cytoplasmic streaming and the intracellular transport of organelles by myosins as early 1774 by the Italian physicist, Bonaventura Corti (Corti, 1774; as cited in Nebenführ and Dixit, 2018). Furthermore, *Chara* cells were used as a model for calcium signaling and plant electrophysiology in the 1970s, owing again to the large cells and researcher’s ability to microinject ions, molecules, and proteins directly into the cytoplasm of individual cells (Williamson, 1975; Williamson and Ashley, 1982). Despite the impressive history of the *Chara* genus in aiding our understanding of the molecular world, specifically calcium-signaling and electrophysiology in plant cells, the calcium-sensing proteins responsible for these processes have not been examined. In an effort to better understand the evolutionary history of the CaM/CML family in plants, we interrogated the *C. braunii* genome to identify putative CaM/CML protein family members. We expressed and purified recombinant CbCaM/CMLs, explored their biochemical properties, and assessed their interaction with a CbCAMTA family member as a representative target protein. We discuss our observations in the context of comparative biochemistry and the evolution of CaM/CMLs in the green lineage.

## Results

### Bioinformatic characterization of the Chara braunii Calmodulin-like protein family

In previous studies, the sequences and characteristics of the CaM and CML proteins of other plant species have been reported (McCormack and Braam, 2003; Zhu et al., 2015; La Verde et al, 2018). Here, using an evolutionarily-conserved isoform of CaM, Arabidopsis AtCaM7, as our query sequence, we identified one *CbCaM* and eight *CbCML* genes encoding non-redundant sequences in the *C. braunii* genome (Nishiyama et al., 2018). The CMLs were named based on their homology to the conserved AtCaM7 sequence (Fig. 1). We used Interpro (Paysam-Lafosse et al., 2022) to predict the basic structural properties and functional domains of the CbCaM/CMLs (Table 1). CbCaM shares 92% protein sequence identity with AtCaM7 and only differs at 11 of 148 residues. These amino acid changes were mostly conserved, aside from the S26C position in EF-hand one and H66P mutation immediately following EF-hand two in AtCaM7 compared to CbCaM (Fig. 1). The predicted molecular weights of the CbCMLs ranged from 16.4 to 21.3 kDa and all are predicted to exhibit acidic pI values ranging from 3.93 to 5.12, typical of the CML protein family (Bender and Snedden, 2013). CbCaM and CbCML3, CbCML5, and CbCML6 are predicted to possess four EF-hands, whereas CbCML2 and CbCML4 are predicted to have three EF-hands, and CbCML1, CbCML7, and CbCML8 are predicted to have 2 EF-hands. The primary structure of CbCML6 has an unusual ∼45 residue acidic, C-terminal extension that is rich in Asp and Asn and is unique among CMLs reported to date. By comparison, CbCML7 possesses a Pro and Ser-rich N-terminal extension that is reminiscent of *Arabidopsis* AtCML48-50, all of which are currently of unknown function. Given that most CML research has been conducted on Arabidopsis isoforms, we performed a phylogenetic analysis of *C. braunii* and included the Arabidopsis CML family. The phylogenetic tree suggests that CbCML1, CbCML2, and CbCML4 are closely related to conserved CaMs, and group with the *Arabidopsis* orthologs AtCML8, AtCML11, AtCML13, and AtCML14. CbCML3 and CbCML8 clustered into the AtCML19 (CEN2) and AtCML20 (CEN) subgroup, while CbCML5 and CbCML7 subdivided into their own group and are most closely related to the Arabidopsis orthologs in subgroup four, AtCML15-AtCML18. None of the CbCML isoforms clustered into the Arabidopsis subgroups six, eight, or nine (Zhu et al., 2015). Interestingly, although CbCML7 possesses the Pro-rich N-terminal extension that is characteristic of the Arabidopsis orthologs AtCML48-AtCML50 within subgroup seven in Arabidopsis, it did not cluster with those Arabidopsis isoforms, whereas CbCML6 which lacks the Pro-rich extension did.

**Figure 1.**
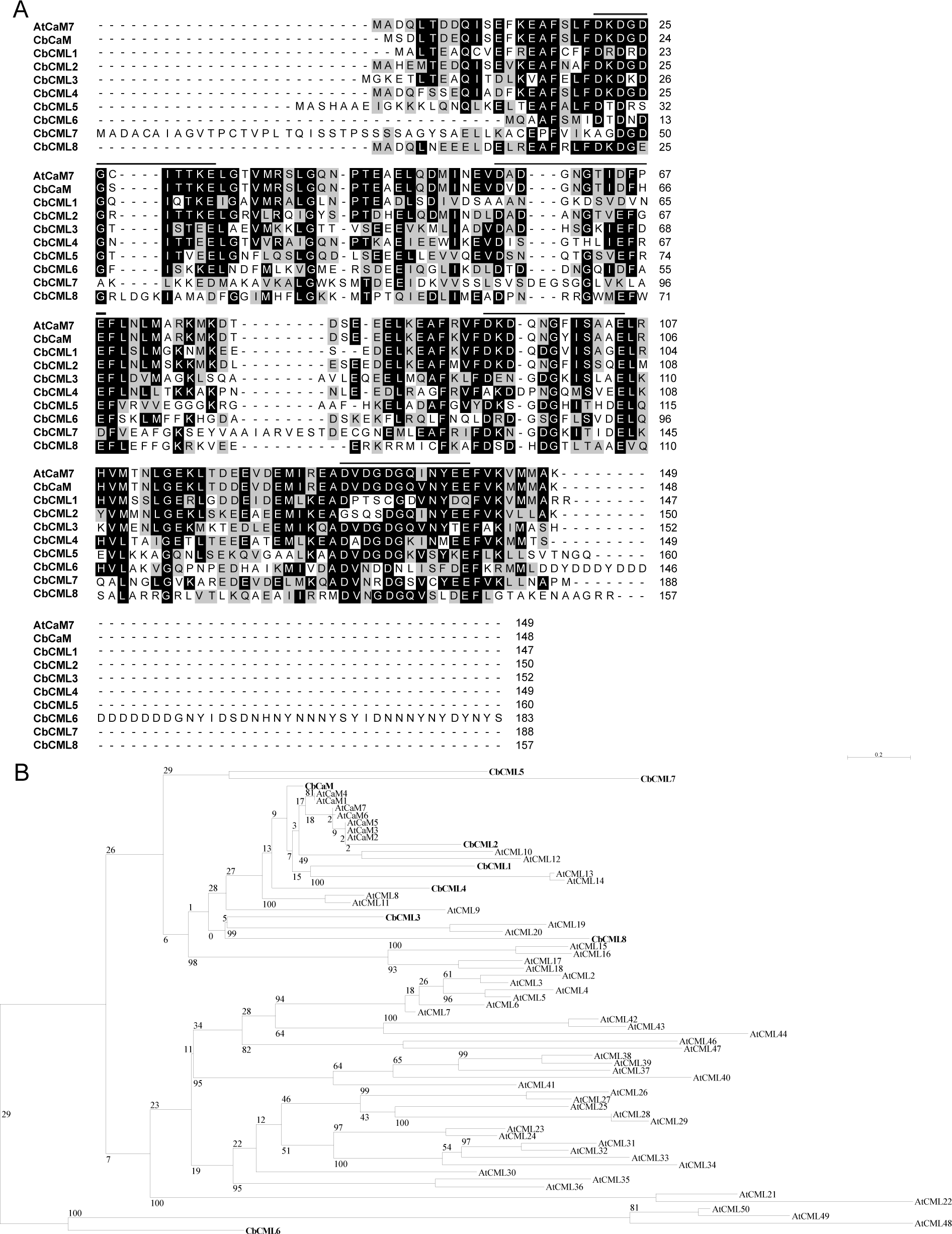
Phylogenetic comparison of *Chara braunii* and *Arabidopsis thaliana* CaM/CMLs. A) Protein sequence alignment of *Chara braunii* CaM and CMLs with a conserved CaM, CaM7, from *Arabidopsis thaliana*. The alignment was generated using Clustal Omega (Sievers and Higgins, 2014) and the figure was produced with BioEdit V7.2.5 (Hall, 1999). Amino acids are shaded black based on >50% conservation, grey if they are functionally conserved changes, and white for non-homologous amino acids. B) Phylogenetic tree of *Chara braunii* and *Arabidopsis thaliana* CaM/CML family members generated with the PhyML algorithm with the Blosum62 scoring matrix and 1000 bootstrap replicates on the SeaView software V5.0 (Guoy et al., 2021).

**Table 1.**
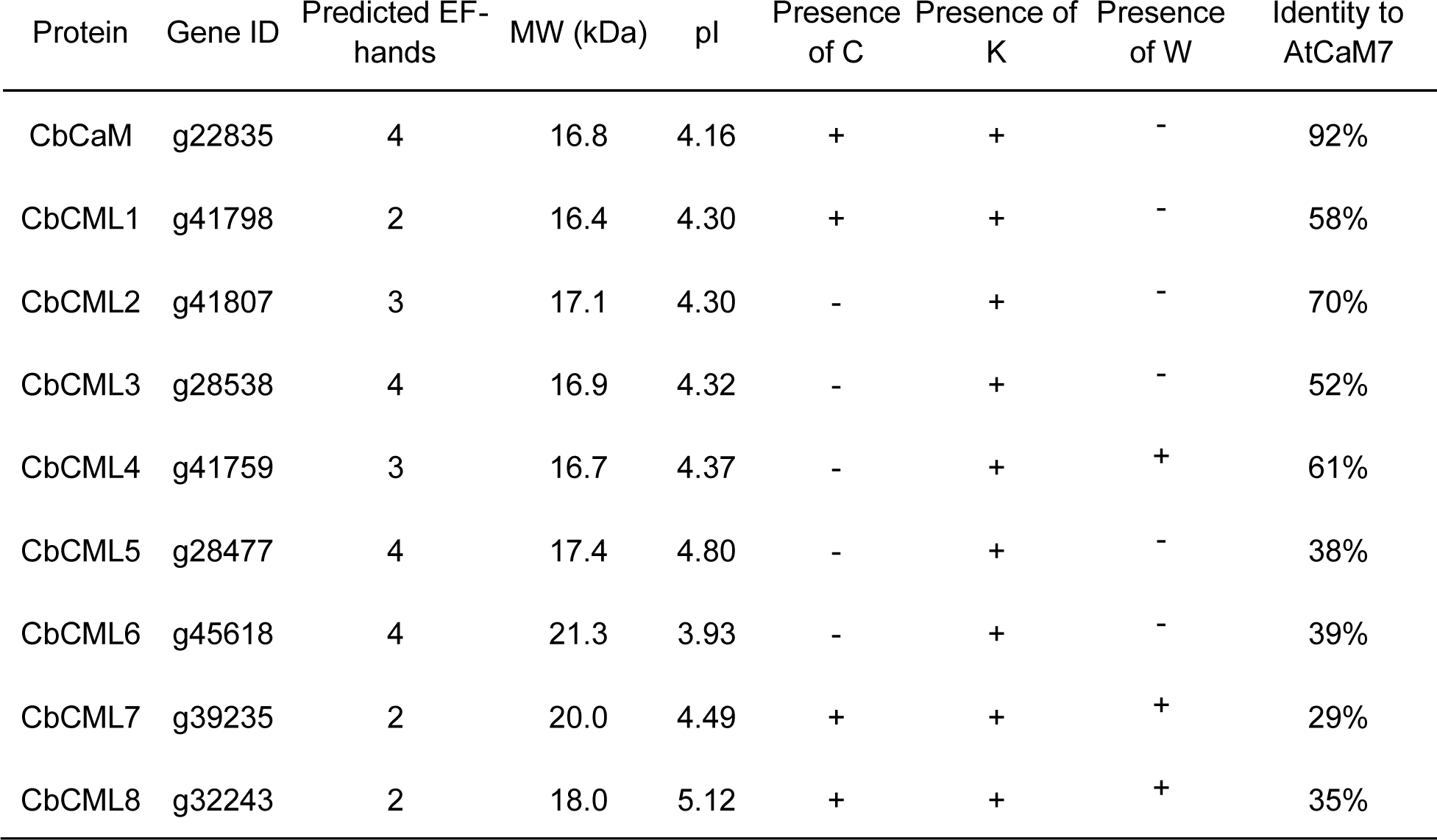
Predicted properties of CaM and CML proteins identified in *Chara braunii*.

We next explored the *in silico* predicted structures of the CbCaM/CML proteins (Fig. 2A). The open-source service ColabFold running AlphaFold2 (Jumper et al., 2021, Miridita et al., 2022) was used to model the tertiary structures of the CbCaM/CML proteins. As depicted in Fig. 2A and 2B, CbCML6 and CbCML7 each possess a C- or N-terminal extension, respectively, that were predicted to form an unstructured loop. The models were then aligned based on their peptide backbones and the root mean squared differences (RMSD) of atom locations were assessed using the RMSD Align tool in the VMD V1.9.3 software (Fig. 2C) (Humphrey et al., 1996). The structures of most CbCMLs were quite similar to each other and to CbCaM (RMSD <10) except for CbCML6 and CbCML7 which had RMSD values closer to 20 compared to the other family members (Fig. 2C). These predicted structural similarities, despite comparatively low sequence homologies in some cases (Supplemental Data Fig. S1), led us to compare the biochemical properties of recombinant CbCMLs *in vitro* to better understand their differences and potential roles as calcium sensors.

**Figure 2.**
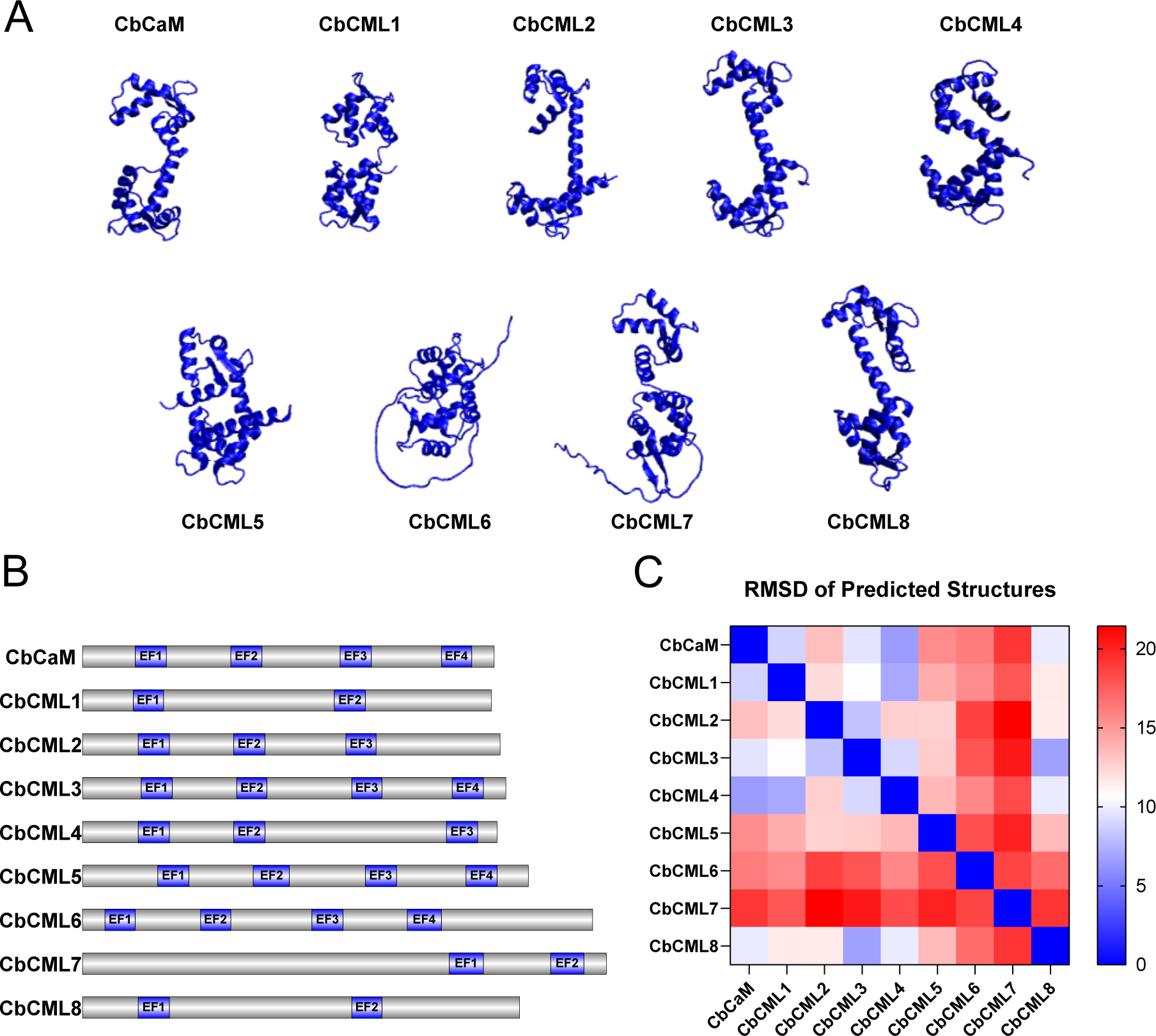
*Chara braunii* CaM/CML protein functional domain organization and predicted structures. A) Predicted protein structures of CbCaM/CMLs generated using ColabFold running AlphaFold2 (Mirdita et al., 2022). Composite image and RMSD structural comparisons were made in VMD V1.9.3. (Humphrey et al., 1996). B) Schematic showing predicted functional domains of the *C. braunii* CaM and CML proteins generated with IBS V2.0 and protein domains predicted using InterPro. C) A heat map of the root mean square difference (RMSD) of atom locations of the CbCaM/CML family members. Scale bar denotes the RMSD values from 0 to 20 where and RMSD = 0 suggests identical atom locations.

### Biochemical characterization of the Chara braunii Calmodulin-like protein family

The cDNA sequences of the *C. braunii* CML family were commercially synthesized, cloned into pET-30a vectors, and the proteins expressed recombinantly. All CMLs were purified via Ni-IMAC resin to near homogeneity except for CbCML8 which was insoluble upon recombinant expression in *E. coli*. Various attempts to stably re-solubolize CbCML8 were unsuccessful and it was not studied further. The main objective of the biochemical analyses was to assess whether the CbCMLs possess properties consistent with their predicted roles as calcium sensors. Structural changes in CbCaM and CbCMLs in response to calcium binding were explored via calcium-dependent electrophoretic mobility shifts, hydrophobicity analysis using ANS (8-anilinonapthalene-1-sulfonic acid) fluorimetry, intrinsic Trp fluorimetry, and circular-dichroism (CD) spectrometry. In addition, we estimated the global calcium affinities of the CbCaM/CMLs using dibromo-BAPTA (5,5-dibromobis-(o-aminophenoxy)ethane N,N,N’,N’-tetra-acetic acid titration analysis.

Increased electrophoretic mobility in the presence of calcium is a characteristic property of CaM and most CMLs studied to date (Garrigos et al., 1991; La Verde et al., 2018). Thus, we performed mobility assays on the purified CbCMLs (Fig. 3). Proteins were heat denatured in SDS-PAGE sample buffer supplemented with either 1 mM CaCl_2_ (Ca^2+^) or 1 mM EGTA (EGTA) and separated by standard SDS-PAGE. CbCaM exhibited the strongest mobility shift, while CbCML3 and CbCML6 also displayed clear increases in mobility in the presence of calcium (Fig. 3). In contrast, CbCML1, CbCML2, and CbCML5 showed only minor shifts in calcium vs EGTA conditions. Interestingly, the electrophoretic mobility of CbCML4 and CbCML7 was unaffected by calcium or EGTA (Fig. 3).

**Figure 3.**
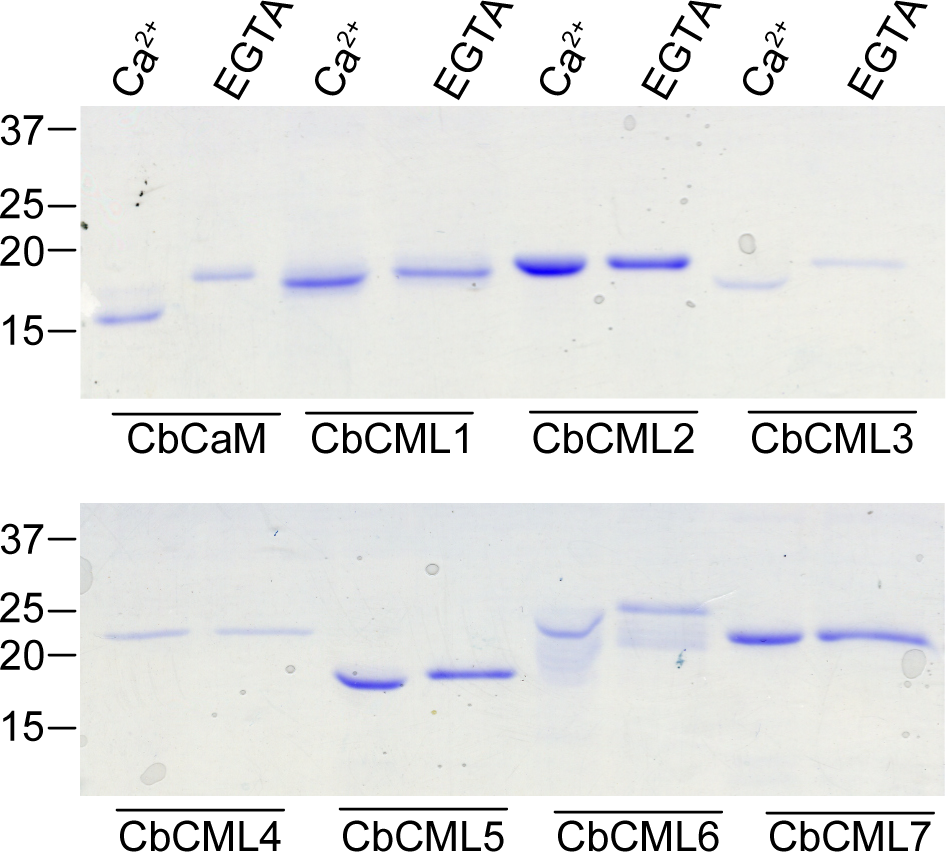
Calcium-dependent electrophoretic mobility shifts of CbCaM and CbCMLs. Recombinant CbCaM/CMLs were run on SDS-PAGE in the presence of 5 mM CaCl_2_ or EGTA as labeled in the figure. The SDS-PAGE gel was stained with Coomassie Brilliant Blue R250 and destained and imaged. Representative images of gels are shown for a minimum of three experimental replicates.

ANS is a fluorescent chemical probe used to assess exposed hydrophobicity and it has been particularly useful in studying calcium-induced conformational changes in CaM and CMLs. These changes are thought to facilitate target interaction in the presence of calcium (Gifford et al., 2007). ANS fluorimetry of recombinant CbCaM/CMLs revealed that CbCaM, CbCML2, CbCML3, and CbCML5, markedly increased their hydrophobic exposure in response to calcium binding, whereas the increase in hydrophobicity of CbCML1 and CbCML6 was not as strong (Fig 4). CbCaM, CbCML2, CbCML3, CbCML5, and CbCML6 exhibited a comparable level of ANS fluorescence in the presence of calcium. Although the ANS fluorescence of CbCML5 was very responsive to calcium, it also displayed a high level of ANS fluorescence in the absence of calcium, suggesting that apo-CbCML5 may have more exposed hydrophobic residues than the other apo-CbCaM/CMLs. Also notable was the observation that although CbCML1 showed a relatively low ANS signal in the absence of calcium, comparable to CbCaM, the increase in hydrophobicity in response to calcium was weaker in comparison to the other calcium-responsive CbCMLs. Also, in the presence of 1 mM MgCl_2_, the fluorescence of CbCML1 was no longer responsive to calcium. In contrast, the hydrophobicity of CbCML4 and CbCML7 appeared to be relatively insensitive to calcium. Given that neither CbCML4 nor CbCML7 exhibited an electrophoretic mobility shift (Fig. 3), yet both possess Trp in their primary sequences, Trp fluorescence was utilized to further assess their ability to bind calcium. The emission spectrum of the Trp in CbCML4 was red-shifted in the presence of calcium which was not observed in the presence of magnesium alone (Fig. 5A). In contrast, the emission spectrum for CbCML7 was unchanged in the presence of calcium or magnesium (Fig. 5B).

**Figure 4.**
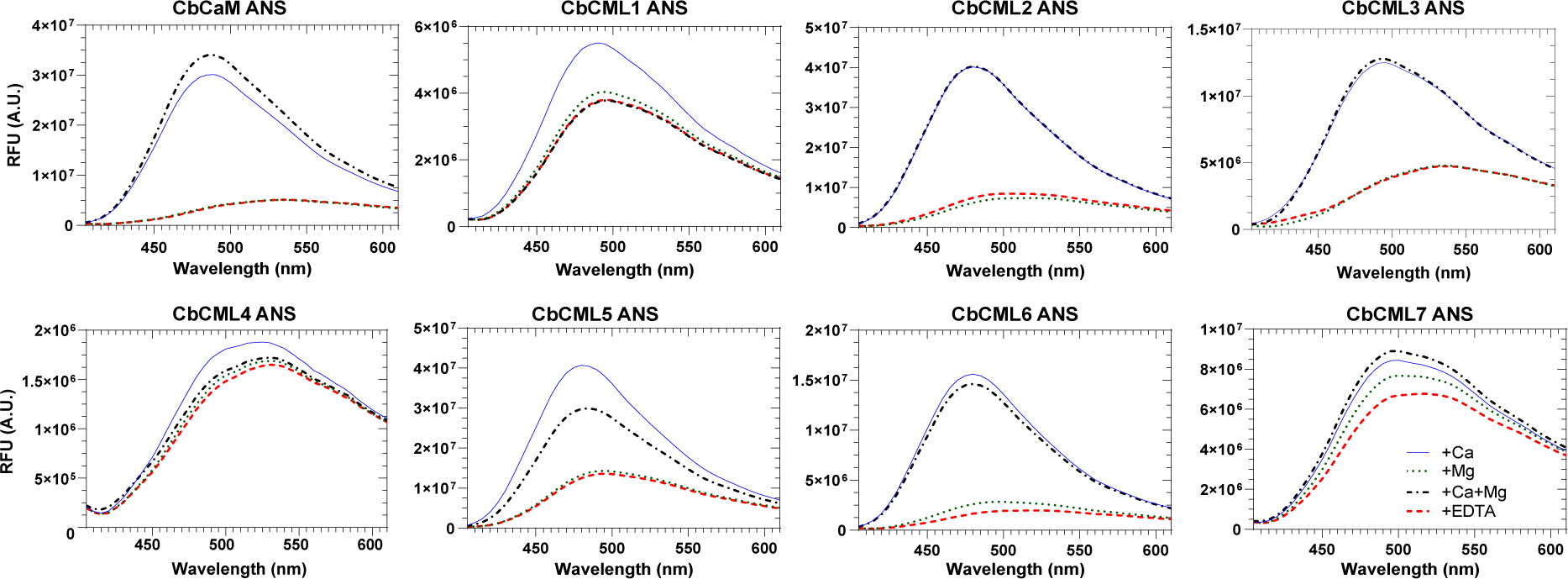
Calcium- and magnesium-induced hydrophobic exposure of CbCaM/CMLs. Recombinant CbCaM/CbCMLs (20 µM) were incubated with 1 mM CaCl_2_ and/or 1 mM MgCl_2_ or 1 mM EGTA and EDTA, as depicted in the figure legend, with the hydrophobicity-sensitive fluorescent compound, ANS (250 µM). ANS fluorescence was measured by exciting at 380 nm and scanning emission from 400-650 nm on the SpectraMax Paradigm (Molecular Devices). Plots show average fluorescence traces of three technical replicates representative of three independent experiments.

**Figure 5.**
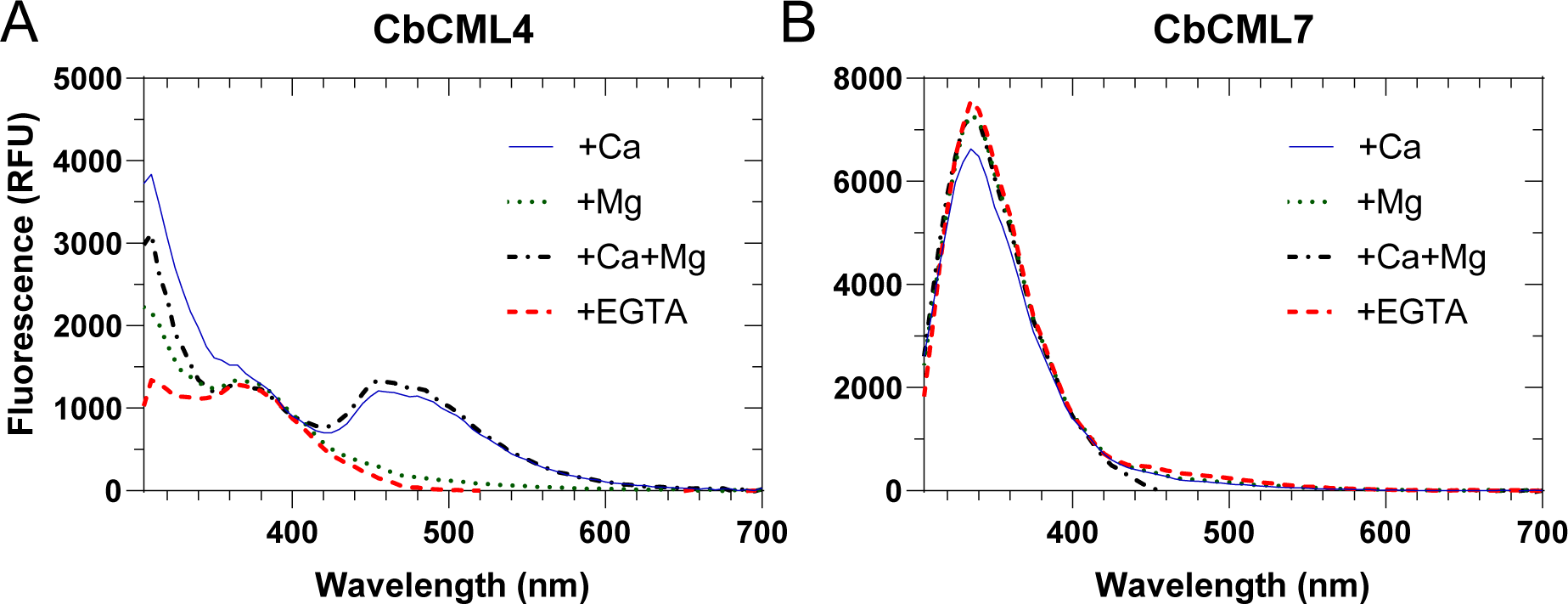
Calcium- and magnesium-induced changes in intrinsic tryptophan fluorescence. Recombinant CbCML4 and CbCML7 (20 µM) were incubated with 1 mM CaCl_2_ and/or 1 mM MgCl_2_ or 1 mM EGTA and EDTA as depicted in the figure legend. Trp fluorescence was measured by excitation at 280 nm and emission was scanned from 300-700 nm on a Synergy-H1 imager (Agilent). Plots show the average fluorescence trace of three technical replicates representative of two independent experiments.

To further characterize the structures of these CbCML family members, far-UV CD spectroscopy was performed. The CD spectrum of CbCaM was similar to that reported for other CaMs, with a strong alpha-helical propensity in the absence of calcium and an alpha-helical gain in the presence of calcium (Zhang et al, 1995; Gifford et al., 2011). CbCML1 displayed relatively weak alpha-helical propensity without calcium. With the addition of calcium, CbCML1 likely undergoes an alpha-helical and beta-sheet gain displayed by the deepening of the trough in the low 200 nm band and a red-shift of the X-axis intercept, respectively (Fig. 6B). CbCML4 and CbCML6 also displayed weak alpha-helical propensity without calcium, however, CbCML4 underwent minimal secondary structural changes with the addition of calcium while CbCML6 displayed a slight beta-sheet gain (Fig. 6E and G). Similarly, CbCML2 displayed little conformational change with the addition of calcium, however, it had a stronger baseline alpha-helical signal compared to CbCML4 and CbCML6 (Fig. 6C). Like CbCML2, CbCML5 also presented a moderate alpha-helical signal, and in the presence of calcium, CbCML5 showed both alpha-helical and beta-sheet gain (Fig. 6F). CbCML3 and CbCML7 have strong alpha-helical propensities and displayed an alpha-helical gain with the addition of calcium (Fig. 6D and H).

**Figure 6.**
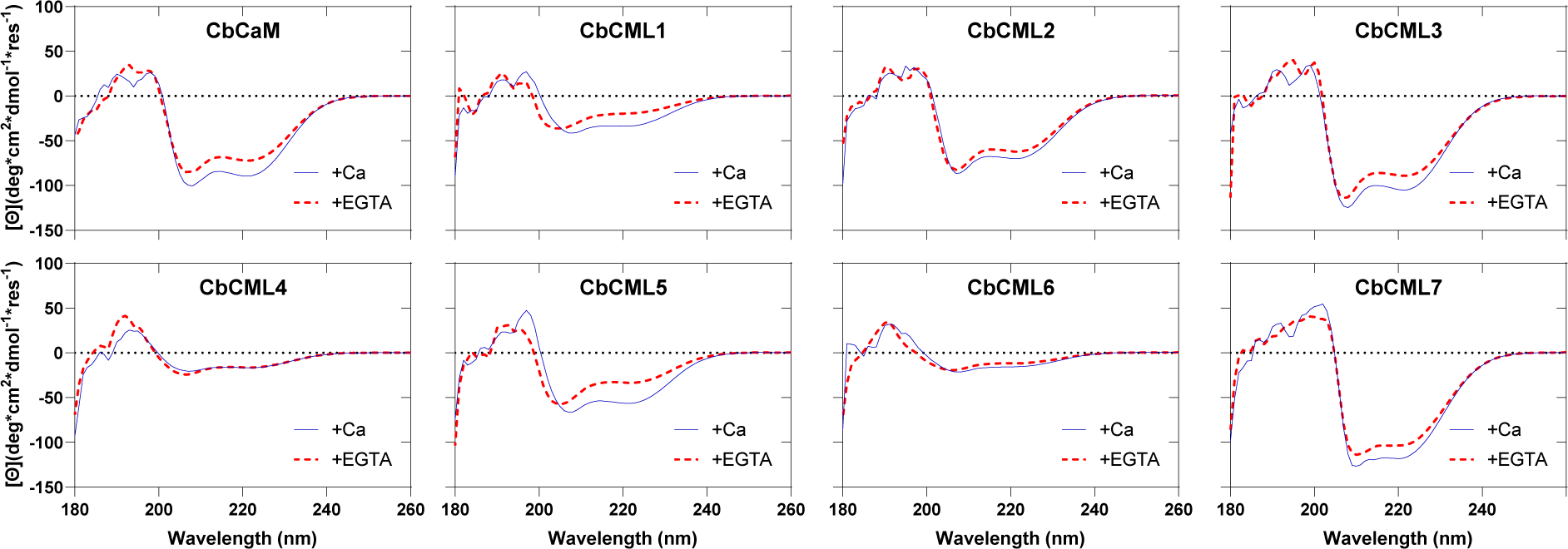
Far-UV circular-dichroism (CD) spectroscopy of CbCaM and the CbCMLs. Recombinant CbCaM/CMLs (25 µM) were analyzed in 5 mM Tris-Cl (pH 7.5) with 25 mM KCl in the presence of either 1 mM EGTA or CaCl_2_ as denoted in the figure. Each spectrum is the average of 5 scans from 180-260 nm using a ChiraScan V100 instrument and is presented in units of molar ellipticity.

### Global calcium affinities of the CbCaM/CMLs

We estimated the global calcium affinities and the biophysical parameters of calcium-binding to CbCaM/CMLs using the calcium titration competition method with the chromophoric chelator dibromo-BAPTA (5,5-dibromobis-(o-aminophenoxy)ethane N,N,N’,N’-tetra-acetic acid) and CaLigator Software (André and Linse, 2002). A representative titration curve for each CbCaM/CML is shown in Figure 7 and the calculated mean +/− SD for the *K*_D_ and Δ*G* of three replicate titrations is presented in Table 2. The global *K*_D_ for calcium among the CbCMLs ranged between 1.10 to 3.06 µM and CbCaM exhibited a *K*_D_ of 2.72 µM which is comparable to that of AtCaM and AtCML isoforms (Dobney et al., 2009; Ogunrinde et al., 2017; La Verde et al., 2018). This method does not allow for stoichiometric analysis, thus the number of functional EF-hands in the respective CbCMLs is based on predictions from primary sequence data. Thermodynamically, Δ*G* values varied from −62 to −144 kJ/mol among the CbCMLs, and this range likely reflects, at least partly, the differences in the number of EF-hands between the isoforms and the extent of conformational changes upon calcium binding.

**Figure 7.**
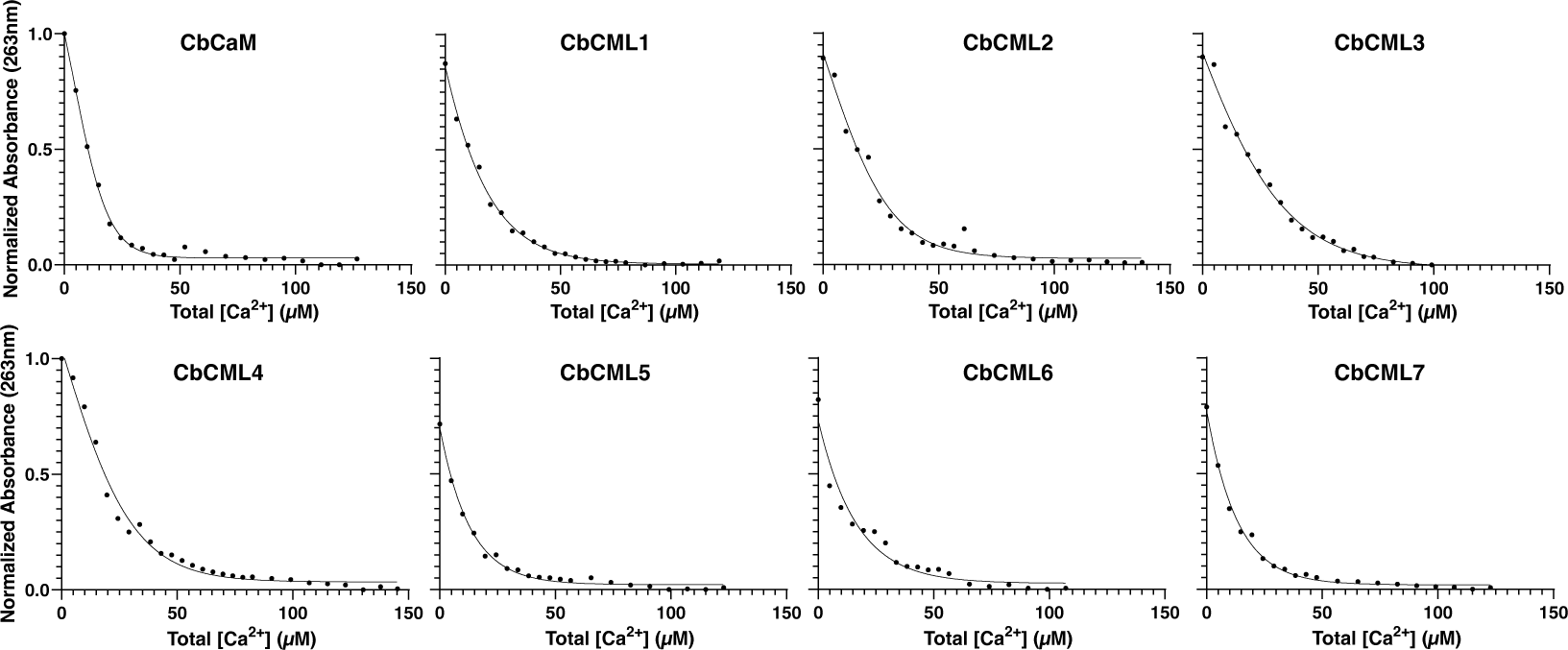
Calcium titration analysis of CbCaM and CbCMLs. Calcium affinities of CbCaM/CMLs were estimated via the competitive titration method with the chromophoric chelator, dibromo-BAPTA. The absorbance of the 28 µM dibromo-BAPTA with 25 µM CbCaM or CbCMLs in solution was measured at 263 nm in a quartz cuvette over a range of CaCl_2_ concentrations until changes in absorbance were negligible. The plots show a representative titration of three experiments. Values for log*K*_A_ and *K*_D_ were estimated via the CaLigator software (Andre and Linse, 2002) using the predicted number of EF-hand domains as reported in Table 1.

**Table 2.** Calcium binding kinetic values of CbCaM and CbCMLs as derived using BAPTA-titration analysis (Fig. 7) and Caligator software as described in Methods.

| Protein | Predicted EF-hands | Global $K_D$ (μM) | $\Delta G$ (kJ/mol) |
| --- | --- | --- | --- |
| CbCaM | 4 | $2.7 \pm 3.2$ | $-120 \pm 7$ |
| CbCML1 | 2 | $1.3 \pm 0.6$ | $-68 \pm 2$ |
| CbCML2 | 3 | $2.6 \pm 2.2$ | $-130 \pm 8$ |
| CbCML3 | 4 | $3.1 \pm 2.8$ | $-130 \pm 3$ |
| CbCML4 | 3 | $1.4 \pm 1.1$ | $-140 \pm 6$ |
| CbCML5 | 4 | $1.7 \pm 2.6$ | $-140 \pm 20$ |
| CbCML6 | 4 | $1.1 \pm 0.6$ | $-140 \pm 7$ |

| Protein | Predicted EF-hands | Global $K_D$<br>(μM) | $\Delta G$<br>(kJ/mol) |
| --- | --- | --- | --- |
| CbCML7 | 2 | 3.6 ± 2.2 | -63 ± 3 |
| CbCML8 | 2 | ND | ND |

### Identification and interaction of CbCAMTA1 with CbCaM/CMLs

As non-catalytic, regulatory proteins, the function of CMLs relies on their interaction with downstream target proteins. Arabidopsis calmodulin-binding transcription activators (CAMTAs) are well-characterized targets of AtCaM and AtCML13 and AtCML14 (Bouche et al., 2002; Yuan et al., 2021; Hau et al., 2023). Thus, we tested whether this interaction is conserved in the *C. braunii* CAMTA and CML orthologs. CAMTAs possess a conventional CaM-binding domain (CaMBD) and multiple CaM-binding IQ (Iso-Gln) domains (Hau et al., 2023). We identified two, non-redundant *CbCAMTA* genes in the *C. braunii* genome and named them based on their sequence similarity to Arabidopsis orthologs. The predicted architecture and domain arrangement of the CbCAMTAs is very similar to that of more recent descendants. For *in planta* binding analysis, the CbCAMTA1 C-terminal domain containing four predicted IQs and a CaMBD was synthesized and cloned into the NLuc vector. The CbCaM/CMLs were cloned into the complimentary CLuc vector and screened for interaction with CbCAMTA1-NLuc using *in planta* split luciferase (SL) assays with AtCML42 as a negative control (Fig. 8B). Among the CbCaM/CMLs tested, only CbCaM, CbCML2, and CbCML4 interacted with the C-terminal domain of CbCAMTA1, and thus their interaction was further characterized. The four putative IQ domains and the putative CaMBD of CbCAMTA1 were tested individually for interaction with CbCaM, CbCML2, and CbCML4 (Fig. 8C-F). CbCaM interacted with the putative IQ4 and CbCML4 interacted with IQ1 and IQ4. Interestingly, CbCML2 did not interact with any of the individual IQ domains tested in the SL assay despite interacting with the full C-terminal region of CbCAMTA1. CbCAMTA1 IQ2 and IQ3 did not interact with any of the CbCMLs tested (Supplemental Data Fig. S2). The isolated CbCAMTA1 CaMBD did not express in the SL assay, thus its interaction with the CbCMLs could not be assessed.

**Figure 8.**
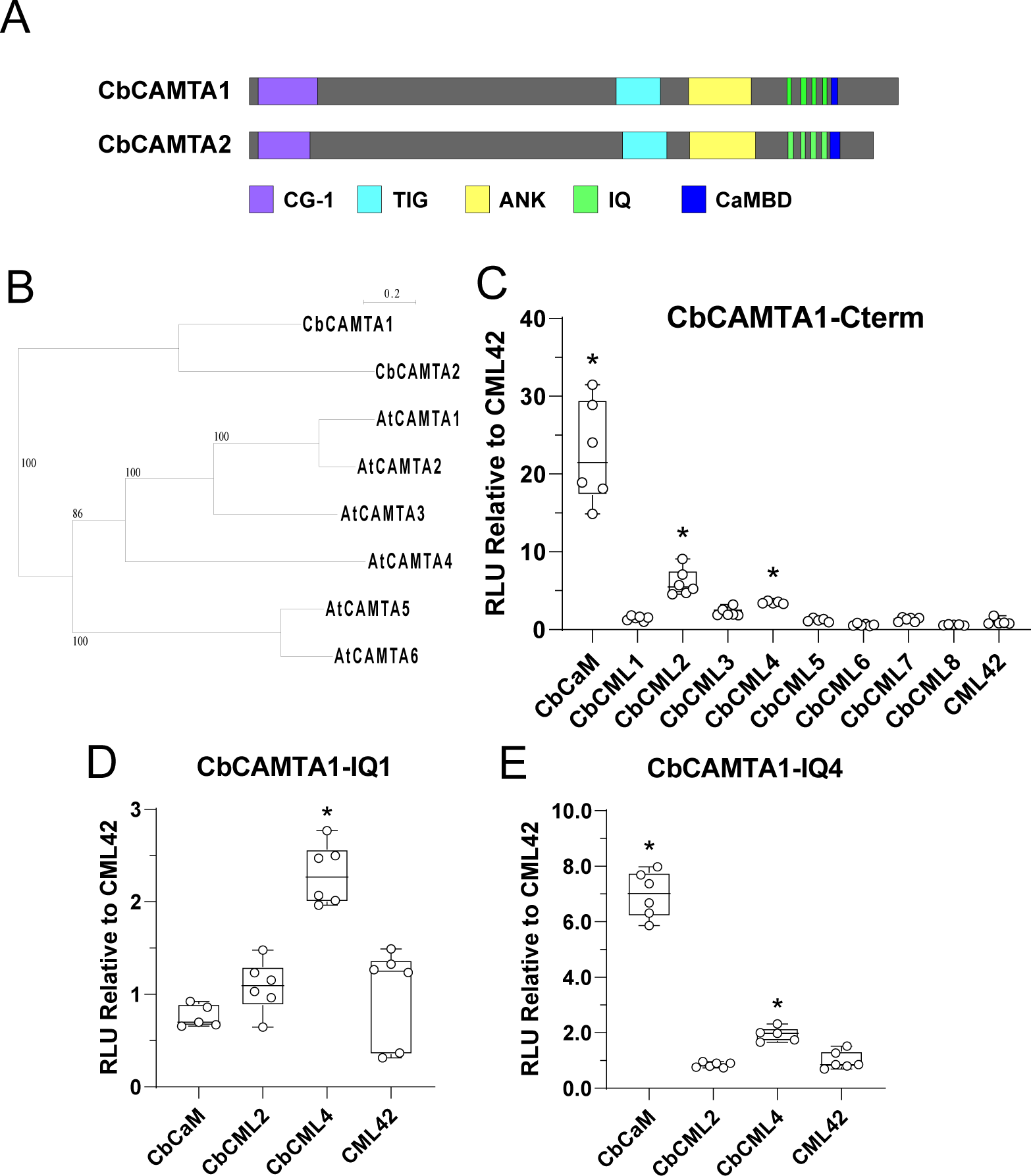
Interaction of CbCaM/CMLs with the C-terminal domain of CbCAMTA1. Two *CbCAMTA* genes were identified in the *C. braunii* genome as being homologous to AtCAMTAs. A) A schematic representation of the predicted domain structure of the CbCAMTAs as predicted by InterPro. The domains depicted are a CG-1 (substrate-specific DNA binding) domain, a nonspecific DNA binding TIG (transcription factor immunoglobulin) domain, ANK (ankyrin) repeat protein-protein interaction domain, multiple IQ (isoleucine glutamine) domains, and a CaMBD (calcium-dependent, conventional CaM binding) domain. B) Phylogenetic tree of CbCAMTA and AtCAMTA proteins generated with the PhyML algorithm with the Blosum62 scoring matrix and 1000 bootstrap replicates on the SeaView software V5.0 (Gouy et al., 2021). C) *In planta* split-luciferase (SL) assays testing interaction of CbCAMTA1-NLuc with each CLuc-CbCaM/CML isoform transiently expressed in *N. benthamiana*. CLuc-AtCML42 was used as a negative control that does not interact with CAMTAs. D, E) The C-terminal domain of CbCAMTA1 was delineated into the individual IQ domains or CaMBDs and tested for interaction with CbCaM, CbCML2, and CbCML4 in SL assays. CbCAMTA1 IQ1 (D) and IQ4 (E) produced positive interaction data while the other IQ domains did not interact (Supplemental Data Fig. S2), and the CaMBD did not express. Split-luciferase data are expressed as a fold difference relative to the negative control bait, AtCML42. Boxes contain each data point for six technical replicates, means are shown by a horizontal bar, the boxed region is the 95% confidence interval, and whiskers extend to maximum and minimum of the data points. Asterisks indicate a significantly higher signal versus CLuc–AtCML42 (one-way ANOVA against AtCML42 with Sidak’s test for multiple comparisons, *P<0.05, **P<0.01, ***P<0.001). Data are representative of at least three independent experiments. RLU, relative light units.

## Discussion

*C. braunii* diverged from higher plants approximately 450 million years ago (Nishiyama et al., 2018), however, the mechanisms for calcium-sensing via the CaM/CML protein family appear to be functionally conserved. This is consistent with the importance of calcium signaling in development and stress response. Exploring the properties of the CaM/CML calcium-sensor family from representative species across the green lineage is an essential step toward understanding how plants have adapted to the challenges of their environments. We identified one Cb*CaM* and eight Cb*CML* genes encoding proteins that share homology with Arabidopsis CaM/CMLs and possess no other predicted functional domains aside from their EF-hands.

The high protein sequence identity (92%) of CbCaM with AtCaM7 confirms it as an evolutionarily conserved CaM isoform. CaM is only found in eukaryotes, and its importance is underscored by the fact that only four other eukaryotic proteins exhibit a greater degree of evolutionary conservation across taxa, namely histones H4 and H3, actin B, and ubiquitin (Ikura and Ames, 2006). Indeed, in comparison to AtCaM7, most of the substitutions in CbCaM occur primarily outside of the EF-hand motifs (Fig. 1). There are two non-conserved variations in CbCaM near EF-hands, S26C within EF1 and H66P following EF2, however, these changes do not appear to have a large impact on calcium affinity as CbCaM displayed a *K*_D_ for calcium in the low µM range that is comparable to that of other CaM orthologs. It is noteworthy that all the calcium affinities we observed for the CbCMLs were in the physiological range, varying from 1.10 to 3.06 µM as estimated via the dibromo-BAPTA titration method (Table 2). Given that Chara cells have been reported to exhibit a resting calcium concentration of approximately 0.22 µM, spiking 30-fold upon excitation to 6.7 µM (Williamson and Ashley, 1982), the calcium affinity values from our analysis suggest that CbCaM/CMLs possess the biochemical ability to function as calcium sensors in *C. braunii*.

In comparing the primary structures of the CbCMLs, we note that the acidic C-terminal extension in CbCML6 is unique among all CMLs/proteins in the NCBI sequence database. The role of this region is unknown, but it does not appear to affect the solubility, folding, or the *in vitro* function of recombinant CbCML6 which displayed conformational changes upon calcium binding and exhibited a low µM *K*_D_ for calcium (Fig. 3, 4, 6, and Table 2). Interestingly, CbCML6 phylogenetically grouped with orthologs AtCML48-50, none of which have an acidic C-terminal extension but, in contrast, all contain a distinct Pro-rich N-terminal region similar to that in CbCML7. Despite this similarity, CbCML7 did not cluster with this subgroup but formed a subgroup with CbCML5. The primary sequence of CbCML5 is more typical of CMLs, but it also has a short N-terminal extension that is predicted to form an alpha-helix, as opposed to an unstructured extension as found in CbCML7 (Fig. 2). Due to its insolubility in *E. coli*, we were unable to investigate the calcium-binding properties of CbCML8. Despite attempts under various conditions, including urea-based denaturation and subsequent renaturation tests, it remains unclear why CbCML8 was insoluble, as the predicted structure is not highly hydrophobic or disordered, especially in comparison to CbCML6 and CbCML7. The primary sequence of CbCML8 is not unusual in any obvious respect, and aside from a four-residue insertion within EF-hand 1, it has high homology to the rest of the family. Nevertheless, recombinant CML insolubility is not uncommon, and in the example of AtCML36, insolubility was due to an unstructured N-terminal region, the truncation of which allowed for purification of soluble AtCML36 (Astengno et al., 2017). Unfortunately, this was not the case for CbCML8, and future studies could attempt to solubilize CbCML8 using solubility protein tags or expression and purification from a eukaryotic system.

Further evidence that CbCaM/CMLs have properties consistent with roles as calcium sensors is the fact that all isoforms tested, with the exception of CbCML4, displayed calcium-dependent conformational changes upon calcium binding (Figs. 4-6). The modest changes in secondary structure in response to calcium, as judged by CDS (Fig. 6), are typical of CaM and CMLs, as they often show more of a rearrangement of helices than a large gain in helical content upon binding calcium (Manning, 1989; Zhang et al., 1995; Dobney et al., 2009). CbCML4 was confirmed to bind calcium by the calcium-dependent red-shifting of the Trp fluorescence spectrum (Fig. 5), likely caused by the proximity of the Trp residue to the EF-hands. However, CbCML4 did not increase hydrophobic exposure (Fig. 4) or show strong changes in secondary structure (Fig. 6) and is reminiscent of Arabidopsis AtCML13 and AtCML14 (Vallone et al., 2016; Teresinski et al., 2023). Further, like AtCML13, the apparent calcium affinity (*K*_D,app_ of 2.3 µM) was measured to be in the low µM range with the dibromo-BAPTA titration assay (Table 2) (Teresinski et al., 2023). It is worth noting that AtCML14 displayed low binding affinity (*K*_D,app_ of 12 µM) for calcium when tested via isothermal titration calorimetry (ITC) instead of the dibromo-BAPTA titration assay (Vallone et al, 2016). This discrepancy between the apparent *K*_D_ measured by dibromo-BAPTA and ITC has been previously noted. For example, the affinity of AtCaM1 for calcium was measured by dibromo-BAPTA and ITC as 13 µM and 20 µM, respectively (Astengno et al., 2016). The differences between their *K*_D_ values are thought to arise from the processes measured, that is the dibromo-BAPTA assay measures competitive calcium binding while ITC measures direct calcium binding and conformational changes of the protein after binding (Astengno et al., 2016). Nevertheless, caution is warranted in extrapolating binding affinities measured under *in vitro* conditions given that ion concentrations, target interactions, or post-translational modifications (PTMs) could alter the calcium-binding properties of CbCaM/CMLs *in vivo*. It will therefore be important in the future to identify the downstream targets and PTMs of CbCaM and CbCMLs.

In general, CaMs and CMLs are thought to affect cell behavior by way of target interaction. We chose a CbCAMTA as a representative putative target of CbCaM/CMLs given that CAMTAs in vascular plants bind CaM and several CMLs (Bouche et al., 2002; Yaun et al., 2022; Hau et al., 2023). Of the Arabidopsis CMLs studied to date, AtCML13 and AtCML14 are unique in that although they show minimal changes in conformation upon calcium-binding, they can interact with high affinity and specificity with the IQ domains of CAMTAs, IQD proteins, and myosins (Teresinski et al., 2023; Symonds et al., 2024b). The CAMTAs of higher plants possess both a conventional CaM-binding domain (CaMBD) and multiple IQ domains that bind CaM and specific CMLs (Hau et al., 2023). Thus, we searched the *C. braunii* genome for predicted CAMTAs and identified two non-redundant sequences and named them CbCAMTA1 and CbCAMTA2. The presence and arrangement of functional domains in the CbCAMTAs are similar to the Arabidopsis orthologs (Fig. 8). We observed the interaction of CbCaM and some CbCMLs with the C-terminal region of CbCAMTA1 *in planta*, suggesting that this property emerged early in the evolution of the green lineage (Fig. 8). In vascular plants, CAMTAs are key transcriptional regulators during biotic and abiotic stress (Iqbal et al., 2020; Yuan et al., 2022), consistent with the importance of calcium sensors under these same conditions. Although aquatic environments are typically more stable than terrestrial ones, the fact that *Chara* CAMTAs interact with CaM/CMLs suggests that the mechanistic links between gene regulation and calcium signaling are consistent across a large evolutionary timeline. The interaction of CbCML2 and CbCML4 with CbCAMTA1 is reminiscent of the binding of AtCML13 and AtCML14 with Arabidopsis CAMTAs. In Arabidopsis, genetic analysis has shown that mutations within or adjacent to CaM and AtCML13/14 binding sites impact CAMTA function, although the specific roles of this interaction remain unclear (Du et al., 2009; Jing et al., 2011; Nie et al., 2012; Kim et al., 2017; Yuan et al., 2021). Given that CAMTAs, including the *C. braunii* isoforms, possess multiple IQ domains, the stoichiometry and specificity of CaM/CML interactions is hypothetically quite complex. Our delineation analyses found that CbCaM interacted with IQ4 of CbCAMTA1, while CbCML4 interacted with IQ1 and IQ4. In contrast, CbCML2 did not interact with any of the delineated domains in the SL assay despite interacting with the full C-terminus and thus CbCML2 may require a larger structure for efficient binding. For example, in Arabidopsis, AtCML13/14 display differential binding to single and tandem (adjacent IQs within ∼12 residues) IQ domains (Teresinski et al., 2023), and a similar phenomenon may occur with CbCML2, however, that remains speculative. Although we did not explore the functional relationship between these protein families, our data imply that the control of CAMTAs via CaM/CMLs and calcium-signaling emerged before the last common ancestor between *C. braunii* and embryophytes. In addition, the CML protein family has been reported to have rapidly expanded following three main evolutionary milestones; multicellularity, terrestrialization, and vascularization (Zhu et al., 2015; Edel et al., 2017). *C. braunii* represents an evolutionarily important species that is multicellular but differentiated from higher plants prior to land colonization (Nishiyama et al., 2018). It would be interesting to investigate in a future study how the expansion of the CML family at each of these evolutionary milestones shaped the CML protein family in plants.

It has been speculated that the expansion of the CaM/CML family in land plants arose because of selective pressure in terrestrial environments where the detection and rapid response to external stimuli would be advantageous (Zhu et al., 2015; Edel et al., 2017). Given that calcium signaling is essential to eukaryotic cell function, we examined the *C. braunii* genome for orthologs of some of the more well-studied calcium signaling components. In addition to the CAMTAs and CaM/CMLs noted above, we found one or more predicted orthologs in *C. braunii* for myosins, IQD proteins, glutamate decarboxylase (GAD), kinesin-like CaM-binding protein (KCBP), CaM-binding protein 60 (CBP60), cyclic-nucleotide gated channels (CNGC), TEOSINTE BRANCHED1/CYCLOIDEA/PCF (TPCs), mechanosensitive ion channel MscS-like (MSL), annexins (ANN), autoinhibited calcium-ATPase (ACAs), calcium exchanger (CAXs), calcium-dependent protein kinases (CDPKs), calcineurin-B like proteins (CBLs) and their interacting protein kinases (CIPKs), and calcium-binding CaM kinase (CCaMK). This list is not exhaustive, but it indicates that the evolution of mechanisms and networks for calcium signaling characterized in vascular plants predates land colonization. Nevertheless, caution is merited when speculating on the roles of these proteins in *Chara* sp. For example, to our knowledge, the only empirical studies showing CaM binding to any *Chara* proteins was the co-purification of a CaM immunoreactive band with *Chara* myosins (Awata et al., 2001) and CbCAMTA1 in the present study. Therefore, it remains possible that while some of these CaM-interacting proteins are involved in calcium signaling in terrestrial plants, the more ancient orthologs in *Chara* may not be. Indeed, the enzyme GAD, which catalyzes the synthesis of gamma-aminobutyric acid (GABA) from glutamate, is a well-studied CaM target in higher plants, and it has been implicated in various stress responses (Fromm, 2020; Shelp et al., 2021). However, the CbGAD orthologs appear to either lack the C-terminal CaMBD that is a hallmark of plant GADs, or have many non-conserved residues in this region, raising the question of whether CbGADs are regulated by CaM (Supplemental Data Fig. S3). In vascular plants, the rapid accumulation of GABA occurs by activation of GAD via Ca^2+^-CaM binding in response to various stimuli including phosphate, salinity, temperature, and drought stress (Snedden et al., 1996; Balfagon et al., 2022; Benidickson et al., 2023). The presence of CbGADs in *C. braunii* indicates that these algae can make GABA, but whether it is integrated into calcium signaling or functions in stress response is not known. Answers to that question and uncovering the roles of the various putative calcium signaling and regulated proteins like fructose bisphosphate aldolase in *C. braunii* will require further analysis (Symonds et al., 2024a).

In conclusion, this study identified and characterized the CaM/CML family of calcium sensors in *C. braunii*. The CbCaM/CMLs show an interesting diversity in their number of EF-hands, calcium-binding properties, and interactions with a representative target, CbCAMTA1. In order to unravel their cellular roles, future studies should empirically explore target identification for the CbCMLs, perhaps using CML orthologs with known targets as a starting point for inquiries. Alternatively, as the techniques of high-throughput protein-protein interaction and proteomics expand, their application to *C. braunii* will help us better understand the evolutionary history of calcium signaling in both charophytes and embryophytes. Similarly, to further the legacy of the *Chara* species as a model for intracellular organelle movement by myosins, the interaction between native CbCaM/CMLs and *C. braunii* myosins should be investigated further. Previous studies on *Chara* myosins reported that the motor domain of myosin CbXI-1 is the fastest known myosin in the biological world, producing actin sliding velocities ten times that of the rabbit fast-twitch muscle fiber and Arabidopsis myosins (Haraguchi et al., 2022). CbCaM alone was unable to support myosin CbXI-1 motility *in vitro* (Haraguchi et al., 2022), and a similar observation was reported for Arabidopsis myosin VIIIs and XIs (Haraguchi et al., 2018). We cautiously speculate that myosin CbXI-1, and possibly other *C. braunii* myosins, may be regulated by CbCML2 and/or CbCML4 along with CbCaM, as based on our biochemical analysis these are likely the functional orthologs of Arabidopsis myosin light chains, AtCML13/14 (Symonds et al., 2024c). Exploring and comparing protein function and cellular processes from deep branching species in the plant lineage provides valuable context into the traits that were selected during evolution. There are a plethora of putative calcium signaling proteins in *Chara* and other evolutionarily important model species that remain unstudied. As such, there is the opportunity to expand the comparative biochemistry and, ultimately, achieve a more holistic picture of how and when the complexity of calcium-based information processing in plants evolved.

## Materials and Methods

### Identification of Chara braunii CaM, CML, and CAMTA genes

The partially annotated *Chara braunii* genome (ORCAE) (Sterk et al., 2012) was searched for proteins/predicted proteins that were homologous to AtCaM7 and contain EF-hand domains (Pfam ID: PF00036, PF13202). Proteins with other functional domains (e.g. kinase domain) were eliminated as CaM/CMLs possess no other functional domains. The CbCaM and CbCML’s amino acid sequences were compared to the conserved AtCaM7 and based on similarity, the CbCMLs were denoted as CbCML1-8 (*Chara braunii* Locus IDs: CbCaM; g22835, CbCML1; g41798, 2; g41807, 3; g28538, 4; g41759, 5; g28477, 6; g45618, 7; g39235, 8; g32243). CbCAMTA genes were identified by sequence similarity to the AtCAMTA3 and CAMTA6 proteins, the CbCAMTAs (CbCAMTA1; g51328, CbCAMTA2; g40845) were named based on their sequence similarities to the AtCAMTA orthologs.

### Cloning and expression of CbCaM, CMLs, and CAMTA1

The cDNA sequences for the full-length *CbCaM* and *CbCMLs* and the sequence corresponding to the C-terminal (AA: 1136-1393) of *CbCAMTA1* with restriction enzyme cut sites were ordered as clonal gene fragments from Twist Bioscience. The full-length cDNAs were digested with KpnI/SalI for *CbCaM/CMLs* or BamHI/SalI for *CbCAMTA1* before being subcloned into pET-30a and C-Luciferase pCAMBIA1300 (CLuc) or pET-28a-SUMO and N-Luciferase pCAMBIA1300 (NLuc), respectively. All vector sequences were verified by Sanger sequencing from The Centre for Applied Genomics (Toronto SickKids Hospital).

Proteins were expressed in *E. coli* strain BL21 CPRIL by growing to OD_600_ = 0.6 and inducing with a final concentration of 0.75 mM IPTG and allowed to express at 37°C overnight. *E. coli* cells were harvested by centrifugation before lysis with a French Press (Glen Mills) at 1100 psi into 50 mM Tris-Cl pH 7.5, 150 mM NaCl (TBS) that was then supplemented with 1 mM PMSF, benzamidine, and DTT (extraction buffer) and the lysate was clarifying through centrifugation again. Proteins were purified by Ni-charged Profinity IMAC resin (BioRad) and purity was analyzed by SDS-PAGE and concentration estimated by the Bradford assay (Bradford, 1976).

### Bioinformatic analysis of CbCaM/CbCMLs

The phylogenetic tree of CbCMLs with AtCMLs was constructed with SeaView 5.0.5 (Gouy et al., 2021) using the PhyML algorithm with 1000 bootstrap replicates and the Blosum62 scoring matrix. Multiple sequence alignment was constructed with ClustalΩ and the figure was made with BioEdit 7.2 (Hall, 1999). Residues were shaded based on conservation with 100% conservation shaded in black and progressively lighter for less similarity. The EF-hand number and locations were predicted by submitting the CbCaM/CML sequences into InterPro and the protein schematic was constructed with Illustrator for Biological Sequences (IBS) 2.0 (Liu et al., 2015).

### ANS and tryptophan fluorimetry

The binding to calcium ions of recombinant CbCaM/CML was monitored using ANS (8-anilinonapthalene-1-sulfonic acid) or intrinsic tryptophan fluorescence emission spectroscopy using an excitation wavelength of 380 nm and emission spectra 430-600 nm or 280 nm and 300-700 nm, respectively (Ogunrinde et al., 2017). The fluorescence emission of 20 μM CbCaM/CML was recorded in a SpectraMax Paradigm (Molecular Devices) using 250 μM ANS in TBS in the presence or absence of calcium and magnesium, as noted in the figure legends. Tryptophan fluorescence was measured for CbCML4 and CbCML7 (the CbCMLs with intrinsic tryptophans) in the presence and absence of calcium and magnesium. Background fluorescence in each buffer was subtracted, and the average trace of three replicates was plotted.

### Circular dichroism (CD) spectroscopy

Far-UV CD spectroscopy of CbCaM/CMLs was performed using a Chirascan CD spectrophotometer (Applied Photophysics, Letterhead, Surry, UK) with cylindrical quartz cuvettes with a pathlength of 1.0 mm. Protein samples were prepared by dialysis overnight in 10 mM Tris-Cl (pH 7.5) and supplemented with CaCl2 or EGTA to a final concentration of 2 mM. Spectra from 10 scans were averaged per assay. The differential absorption of left and right circularly polarized light is presented as the molar ellipticity (θ).

### Di-bromo-BAPTA absorbance titration

Calcium binding affinity to CbCaM/CMLs was assessed using titration analysis as previously described (Pedretti et al., 2020), based on the competition for calcium between the protein and the chromophoric chelator 5,5′-Br_2_-BAPTA [5,5′-dibromobis-(o-aminophenoxy)ethane N,N,N’,N’-tetra-acetic acid] (Biotium). All solutions were decalcified using Chelex-100 resin (Sigma) as per the manufacturer’s instructions.

Calcium titrations, from 0 to 100 μM, were performed by combining a similar concentration of CbCaM/CML and chelator in 5 mM Tris-HCl, 150 mM KCl, pH 7.5. The chelator concentration was 28 μM and the initial calcium concentration was ∼1-5 μM. The absorbance at 263 nm (λ_max_ for the calcium free chelator) was measured as 5 μL aliquots of 1 mM CaCl_2_ were added until no significant changes in absorbance were detected. CaLigator software (André and Linse 2002) was used for data analysis, using a model with a chelator of known calcium affinity (*K*_D_ (5,5′-Br_2_-BAPTA) of 1 μM) and 2-4 macroscopic calcium binding constants for the protein (log K_A_). The apparent affinity value (*K*_D,app_) was determined from the average of the logarithms of the four macroscopic binding constants (*K*_D,app_ = 10^−((logK1+logK2+logK3+logK4)/4)^) as described by André and Linse (2002).

### Split-luciferase interaction assay

*Nicotiana benthamiana* seeds were oversown into Sungro sunshine mix #1 and placed in a Conviron MTR30 growth chamber for 14 days (∼200 µmol/m^2^/s light intensity from RB LEDs, short day: 12 hr light and dark cycle, 23°C, and 70% relative humidity). These *N. benthamiana* seedlings were transplanted to individual pots and grown for another four weeks before infiltration. Agrobacterium harboring either NLuc-CAMTA or CLuc-CaM/CML were infiltrated into *N. benthamiana* leaves as previously done (Teresinski et al., 2023). Luciferase activity was measured 4 days post-infiltration with six technical replicates at least three independent times. All RLU values were made relative to the signal obtained with CLuc-AtCML42 as the negative control.

### Statistical analysis

All data and graphs were analyzed with GraphPad Prism 10 unless otherwise stated.

## Data Availability

All data supporting the findings of this study are available in this paper and in its supplementary data published online.

## Author Contributions

All authors contributed to experimental planning and design. KS, UW, LD, and WS performed the experiments and analyses; all authors contributed to the writing, editing and approval of the manuscript.

## Supplemental Figure Legends

**Supplemental Figure S1.**
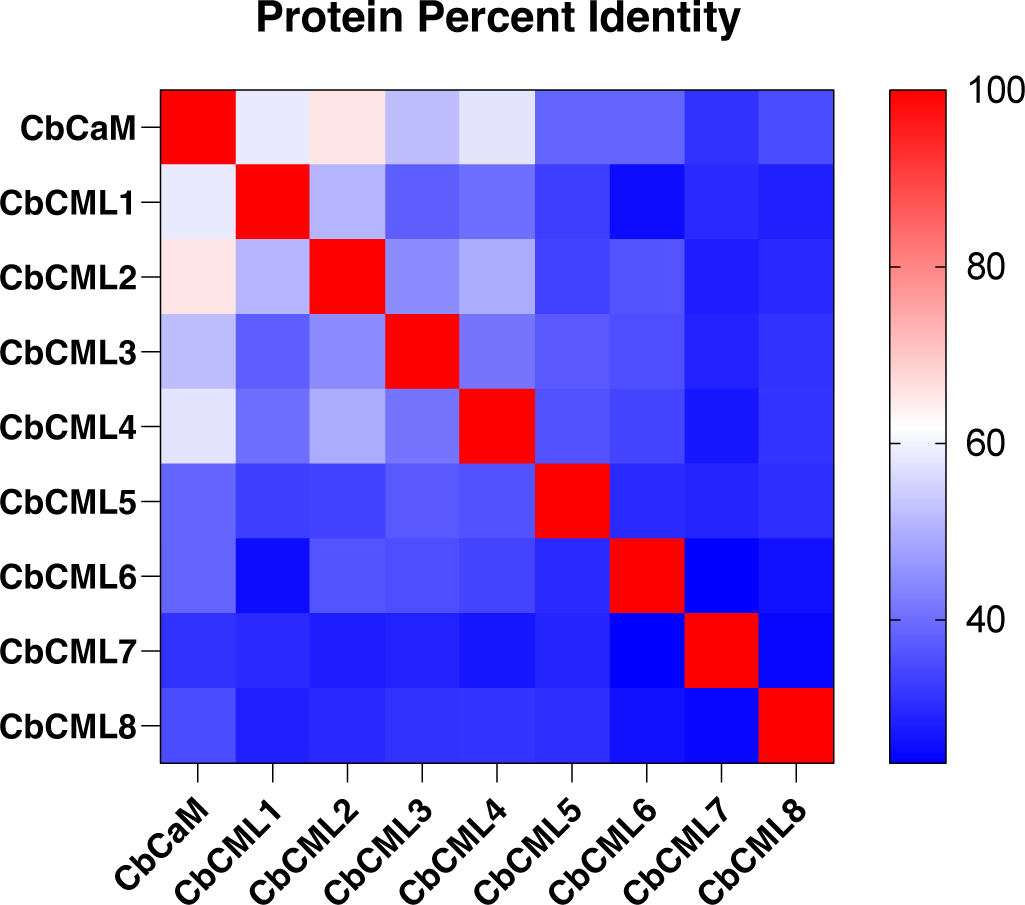
A heat map of the percent identities of the CbCaM/CML family members. Data was generated by Clustal Omega and the figure was made in Graphpad Prism 10. Scale bar denotes the percent identity values from 0 to 100 percent.

**Supplemental Figure S2.**
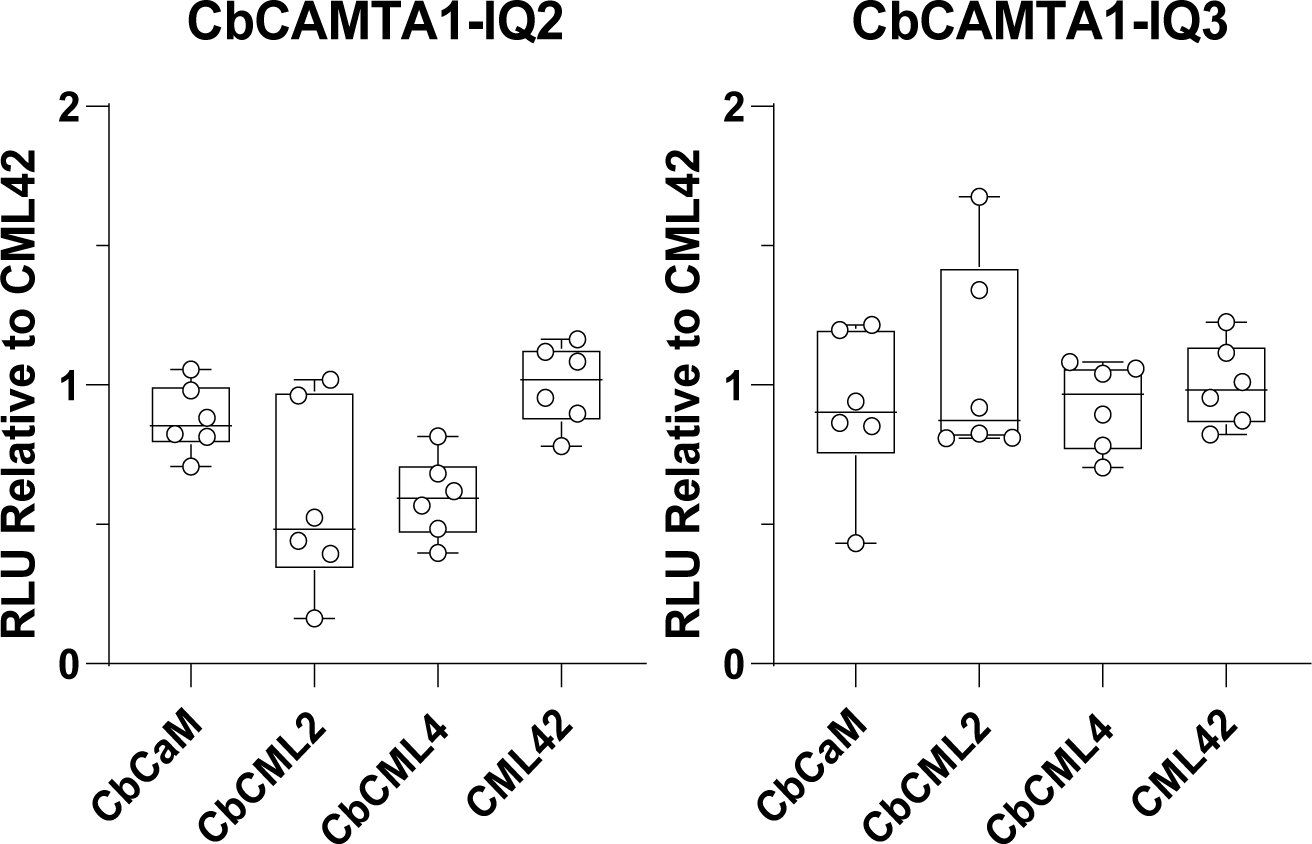
*In planta* split-luciferase (SL) assays testing interaction of CbCAMTA1-NLuc IQ2 and IQ3 with CLuc-CbCaM/CML2/4 isoforms transiently expressed in *N. benthamiana*. CLuc-AtCML42 was used as a negative control that does not interact with CAMTAs.

**Supplemental Figure S3.**
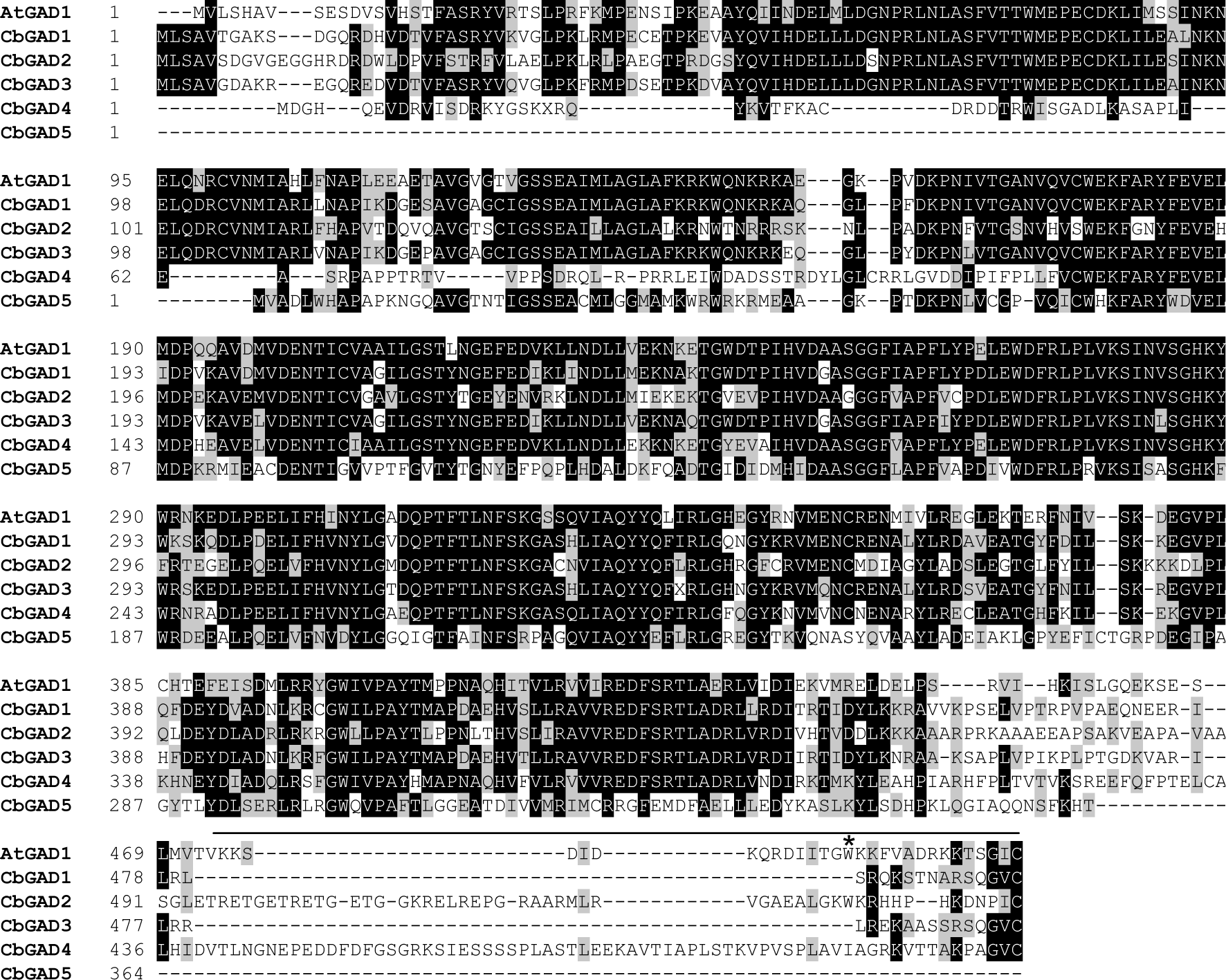
Protein sequence alignment of *Chara braunii* glutamate decarboxylase (GAD) with GAD1 from *Arabidopsis thaliana*. The alignment was generated using Clustal Omega (Sievers and Higgins, 2014) and the figure was produced with BioEdit V7.2.5 (Hall, 1999). Amino acids are shaded black based on >50% conservation, grey if they are functionally conserved changes, and white for non-homologous amino acids. Black bar represents the location of the CaMBD sequence of AtGAD1.

